# Exposure to Wood Smoke is Associated with Increased Risk of Asthma and Respiratory Symptoms in a Honduran Population

**DOI:** 10.1101/085407

**Authors:** Bo Hyun Cho, Elizabeth Castellanos, Elizabeth Nguyen, Sam Oh, Neeta Thakur, Jaime Tarsi, Tammy Koch, Erika Flores de Boquin, Alberto Valladares, John Balmes, Esteban Burchard, Mario Castro, Joshua Galanter

## Abstract

**Background:** Exposure to environmental pollutants has been shown to be associated with asthma, but few studies have evaluated the effect of wood smoke on asthma and disease severity in a developing country, where use of stoves powered by solid fuels is a common practice.

**Objective:** In a population in Olancho, Honduras, we evaluated the association between cooking fuel, stove type and asthma. We also evaluated the effects of these factors on asthma symptoms, lung function, and atopy.

**Methods:** Participants with physician-diagnosed asthma (n = 597) and controls without asthma (n = 429) were recruited from the Olancho province in Honduras. Participants were interviewed using a questionnaire and their baseline pulmonary function was measured using spirometry.

**Results:** The prevalence of use of wood as a cooking fuel was 66.9% in the study population, of which 42.1% of participants used wood as their only fuel. Use of wood as a cooking fuel was more prevalent among households with lower income, lower maternal education, and less urbanization. The prevalence of use of an open wood stove as the primary cooking stove among participants with asthma was 6.2% higher (95% CI 0.8 – 11.7%, p = .02) than among healthy controls. In a multiple logistic regression model, we identified a significant association between use of an open wood stove and asthma (OR = 1.80, 95% CI = 1.17 - 2.78, p = 0.007), compared to the referent (electric) stove category. Among participants with asthma, we identified a significant association between use of wood as cooking fuel and increased daytime respiratory symptoms (OR = 1.46, CI: 1.01 – 2.58, p = 0.046) and nocturnal symptoms (OR = 2.51, CI: 1.04 - 2.62, p = 0.04), though not with pulmonary function. Among control participants without asthma, use of wood as cooking fuel was associated with atopy (OR = 1.94, CI = 1.14 – 3.33, p = 0.015) and cough (OR = 2.22, CI = 1.09 – 4.88, p = 0.04).

**Conclusions:** Use of an open wood stove for cooking in a developing country appears to be a significant risk factor for asthma and respiratory symptoms. Exposure to wood smoke may play a role in atopic sensitization and respiratory symptoms, leading to the development of obstructive lung disease in susceptible individuals.

## Introduction

Asthma is a chronic illness that affects over 200 million people worldwide, and is the 14th most important disorder in terms of global years lived with disability^1^. The increase in prevalence of asthma in the past few decades has been associated with increase in atopic sensitization and urbanization. What has once been considered to be a problem of the developed world is now increasingly becoming a burden in the developing world as well. Due to the combination of an increasing burden of asthma and limited availability of economic and healthcare resources, most asthma-related deaths now occur in low and lower middle income countries^2^.

Worldwide, about 40% of all households and 90% of rural households combust solid fuels for cooking or heating^3–5^. Solid fuels, which include biomass such as dung, charcoal, wood, or crop residues, as well as coal, release air pollutants such as carbon monoxide, benzene, formaldehyde, free radicals, and particulate matter including PM_10_ and PM _2.5_^6^. Particulate levels in houses with open fires are 300 to 3000 μg/m^3^ and may rise to 10,000 μg/m^3^ during cooking. In contrast, the WHO recommends a 24-hour mean PM_2.5_ of < 25 μg/m^3^^7^. This has resulted in an estimated 2 million premature deaths annually attributed to exposure to household air pollution^4^. Household air pollution from solid fuel combustion has been shown to be associated with COPD^8,9^ and lung cancer in adults^10,11^ and lower respiratory infections in children^12,13^, but there is limited evidence that shows the effect of exposure to household air pollution on asthma.

Studies have shown that both asthma prevalence and asthma exacerbations amongst individuals with pre-existing asthma are associated with outdoor pollution^14,15^ and childhood exposure to secondhand smoke from tobacco is a risk factor for development of asthma and allergic diseases^15^. Because household air pollution from solid fuel combustion bears an even greater concentration of air pollutants than outdoor pollution or secondhand tobacco smoke, we would expect to observe a significant association between exposure to household air pollution and asthma. A few studies have shown increased risk of asthma with household air pollution from solid fuel use, and a meta-analysis of four studies showed that exposure to household air pollution doubled the risk of developing asthma in children^16^, though those findings were not supported in a separate meta-analysis^17^. A recent analysis of phase three of the International Study of Asthma and Allergies in Childhood (ISAAC) found that the use of open fires (compared to electric or gas stoves) for cooking was associated with an increased risk of symptoms of asthma and reported asthma^18^.

The Honduran Latino Asthma (HOLA) study seeks to investigate the effect of wood smoke exposure on asthma in Honduras, the sixth least developed country in Latin America. We examined how different stove types and cooking fuels were associated with physician diagnosis of asthma, symptoms of asthma, lung function and other atopic symptoms in the towns of Juticalpa (population 130,000) and Catacamas (pop. 122,000), and nearby semi-urban and rural areas in Olancho, Honduras.

## Materials and Methods

### Study Design and Study Population

The Honduran Latino Asthma (HOLA) study is an ongoing case-control study of asthma in children and young adults (ages 5 – 40). Participants were recruited from the towns of Juticalpa and Catacamas, and nearby semi-urban and rural areas in the Olancho department in Honduras. Participants with asthma were recruited from the Hospital San Francisco in Juticalpa and Hospital Santo Hermano Pedro in Catacamas. Healthy controls were drawn from the local communities, with attempts to frequency match based on age, sex, socioeconomic status, and geography. The study protocol was approved by the Committee on Human Research at the University of California, San Francisco and the Institutional Review Board at the Washington University, St. Louis, as well as the Committee of Investigations at the Hospital Católica de Honduras. Written informed consent was obtained from all participants. For participants under the age of 18, parental consent as well as age-appropriate participant assent was obtained.

Asthma cases were defined as participants 1) with a physician diagnosis of asthma, including those newly diagnosed and 2) who were taking medications to control asthma or 3) who reported 2 or more of asthma symptoms (wheezing, dyspnea, nocturnal awakening, or cough not associated with a respiratory tract infection) in the 12 months prior to enrollment. Controls without asthma were defined as participants who did not meet any of the above inclusion criteria for asthmatic participants. All potential participants were excluded if they 1) had a diagnosis of any other cardiac or pulmonary disease or 2) had been active smokers in the previous year or had a greater than 5-pack-year history of smoking or 3) were unable to complete any part of the study.

All participants were given a detailed questionnaire that was formulated based on large pediatric and adult asthma cohort studies including the National Cooperative Inner-City Asthma study^19^, the Fresno Asthmatic Children’s Environment Study (FACES)^20^ the Oakland Kicks Asthma Study, the GALA II study^21,22^, the Chronic Respiratory Effects of Early Childhood Exposure to Respirable Particulate Matter (CRECER) study^23,24^, and the ISAAC study^25,26^, and from validated instruments such as the Asthma Control Test^27,28^. The questionnaire evaluated the characteristics and the extent of wood smoke exposure, including cooking fuel use and stove type, as well as other environmental exposures, socioeconomic information, and early life health and environmental information. Spirometry was performed according to ATS/ERS guidelines at the time of enrollment^29^. Because there are no spirometric reference equations specifically for Hondurans, predicted values were generated using Hankinson equations for Mexican Americans^30^ and both raw spirometric values, adjusted for age, sex, height, and height squared and percent predicted values were analyzed.

### Environment and Stove Types

Cooking fuel and stove type were assessed by self-report. The majority of participants reported using either wood, propane, or electricity, or a combination of the above as a cooking fuel. For the purposes of analysis, a three-factor categorical variable of fuel use was generated, assigning participants to households based on whether wood was the sole cooking fuel source, was used in combination with another fuel source, or another fuel source (propane or electricity) was the sole fuel source. Participants were also asked about their primary stove type; options included open wood stove, improved wood stove with ventilation or chimney, propane stove, electric stove, or other stove type.

### Data Analysis and Regression Model Adjustments

Logistic regression models were used to model binary outcomes such as asthma status, presence of atopy, and presence of cough. Ordinal logistic regression was used to model ordered outcomes, such as daytime symptom frequency, nocturnal symptom frequency, degree of limitations, rescue inhaler use frequency, and level of asthma control. Linear regression was used to model continuous outcomes such as pulmonary function measurements. All models were adjusted for appropriate covariates, including age, sex, measures of socioeconomic status including annual household income and maternal education, urbanization (rural, small town, or city), and presence of household smokers. Measures of asthma severity, including lung function in participants with asthma, were also adjusted for use of controller medications such as inhaled corticosteroids. All analyses were done in the R statistical language version 3.3.0^31^.

## Results

Baseline characteristics of the HOLA study population are shown in Table 1. Asthma cases and healthy controls had a similar age and sex distribution. Although a substantial majority of both groups described themselves as “Mestizo”, more controls than cases self-described as European and Native American ethnicity. Cases and controls had a similar annual income. As expected, participants with asthma had a significantly lower FEV_1_ as percent of predicted and a lower FEV_1_ /FVC ratio than healthy controls.

**Table 1:** Baseline characteristics of HOLA participants. For continuous variables, median (25:75 interval) is given. The p-value is determined with a Fisher’s exact test for categorical variables and a t-test for continuous variables. The approximate current value of the Lempira is 21 HNL to 1 USD.

|  | Asthma Cases | Healthy Controls | p |
| --- | --- | --- | --- |
| Number | 597 | 429 |  |
| Age | 20 (13: 29) | 21 (19: 24) | 0.5 |
| Sex (% Male) | 223 (37.3%) | 169 (39.3%) | 0.4 |
| Race/Ethnicity* |  |  | 0.06 |
| European/White | 5 (0.8%) | 10 (2.3%) |  |
| Native American | 4 (0.7%) | 9 (2.1%) |  |
| African/Black | 4 (0.7%) | 3 (0.7%) |  |
| Mestizo (Mixed) | 578 (96.8%) | 400 (93.2%) |  |
| Other† | 6 (1%) | 7 (1.6%) |  |
| Annual Income (Lempiras) | 6,000 (3,000: 14,100) | 8,000 (4,000: 23,000) | 0.2 |
| Pulmonary Function |  |  |  |
| FEV1 (% predicted) | 91.0 (81.6: 101.9) | 92.9 (85.8: 100.2) | 0.04 |
| FVC (% predicted) | 92.5 (82.7: 102.1) | 90.9 (82.7: 98.0) | 0.1 |
| FEV1/FVC | 0.86 (0.80: 0.91) | 0.89 (0.85: 0.94) | $7.3 \times 10^{-11}$ |

\* Concepts of Race/Ethnicity in Honduras differ from those in the United States; most participants self-describe as "Mestizo" or mixed ancestry. Under US census guidelines, most/all participants would be considered of Latino ethnicity.
† Most common "other" response was "Honduran"

**Table 2:** Association between woodsmoke exposure and asthma.

\*Odds ratios calculated by comparing the stated stove type in comparison to the reference stove (electric stove).
| **Risk Factor** | **Asthma**<br>**(Odds Ratio, 95% CI)** | **p** |
| --- | --- | --- |
| Use of open wood stove as main cooking source* | 1.8 (1.2-2.8) | 0.007 |
| Use of ventilated wood stove as main cooking source* | 1.12 (0.7-1.8) | 0.6 |
| Use of open wood stove compared to all other stove types | 1.55 (1.08-2.22) | 0.02 |
| Use of wood as fuel | 1.18 (0.8-1.72) | 0.4 |

**Table 3:** Use of wood as cooking fuel as a risk factor for various symptoms among asthma cases and controls. Results show significant association of use of wood as fuel with increased asthma symptoms in cases and with non-asthmatic allergic symptoms in controls.

| **Symptoms** | **Asthma Cases**<br>**(OR, 95% CI)** | **Healthy Controls**<br>**(OR, 95% CI)** | **p** |
| --- | --- | --- | --- |
| Daytime symptom frequency | 1.46 (1.01-2.58) | - | 0.046 |
| Nocturnal symptom frequency | 2.51 (1.04-2.62) | - | 0.04 |
| Atopy* | - | 1.94 (1.14-3.33) | 0.015 |
| Chronic cough | - | 2.22 (1.09-4.88) | 0.036 |

Overall, wood was used as a cooking fuel in 66.9% of participants’ households, and as the only cooking fuel in 42.1%. Annual income was a strong predictor of use of wood as a fuel (Figure 1A); participants who use wood as their only cooking fuel had an annual income of 4,000 HNL (Honduran Lempira, approximately $190) (25:75% 2,000: 10,000 HNL), compared to 6,000 HNL (3,550: 10,000 HNL) in participants who used both wood and another source of fuel, and 10,000 HNL (5,000: 16,000 HNL) in participants who only used another source of fuel (Kruskal-Wallis rank sum test p-value 1 × 10^−12^). We also found that maternal education (Figure 1B) and degree of urbanization (Figure 1C) were inversely associated with use of wood as cooking fuel (χ^2^ p-value 5 × 10^−15^ and 8 × 10^−14^, respectively). We found similar associations between primary stove type and income, with participants who used propane (median income 10,000 HNL, 25:75% 5,000: 24,000 HNL) or electric stoves (10,000; 5,000: 20,000 HNL) having higher incomes than those using open wood stoves (5,000; 2,350: 13,500 HNL) or improved wood stoves (4,000; 2,000: 7,000 HNL). Because these were potential confounders of the association between cooking fuel, primary stove type and asthma, adjusted regression models included annual income, degree of urbanization, and maternal education as covariates in multivariate regressions.

**Figure 1.**
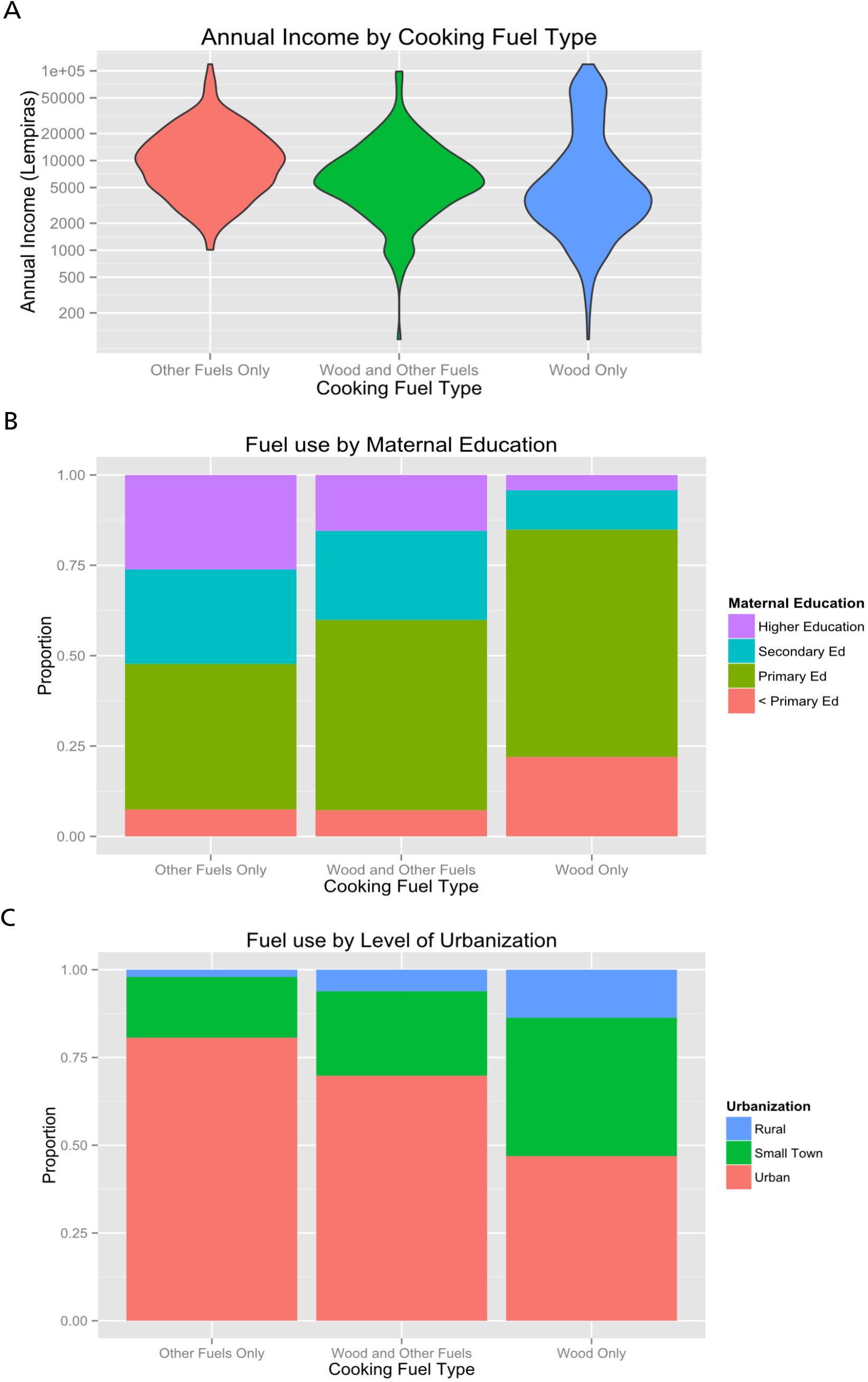
Factors associated with fuel type use. A) Violin plot showing annual income (in HNL; $1 USD ≈ 21 HNL) by wood stove use. B) Bar chart showing maternal education by wood stove use. C) Bar chart showing degree of urbanization by wood stove use. Wood stove use is associated with lower annual income, lower maternal education, and lower degree of urbanization.

We found differences in primary cook stoves but not fuel type between participants with asthma and controls (Figure 2). Specifically, among participants with asthma, 26.8% used an open wood stove as their primary stove for cooking, 18.6% of subjects used improved wood stoves with ventilation, 26.5% of subjects used propane stoves, and 26.8% subjects used electric stoves as their main stoves, while among healthy controls, 20.6% of subjects used open wood stoves, 21.3% of subjects used improved wood stoves with ventilation, 27% of subjects used propane gas stoves, and 29.6% of subjects used electric stoves as their main stoves (Figure 2B). In the unadjusted analysis, the proportion of participants with asthma who used open wood stove as their primary stove was 6.2% higher (95% CI 0.7% to 11.7%, p = 0.03) than the proportion of controls using primarily open wood stoves. In contrast, similar proportions of patients with asthma and controls used wood as their only fuel source (43.3% vs. 40.4%, difference 2.9%, 95% CI -3.4% to 9.2%, p = 0.4) or as one of multiple fuel types (24.3% vs. 25.4%, difference -1.1%, 95% CI -6.7% to 4.4%, p = 0.7).

**Figure 2.**
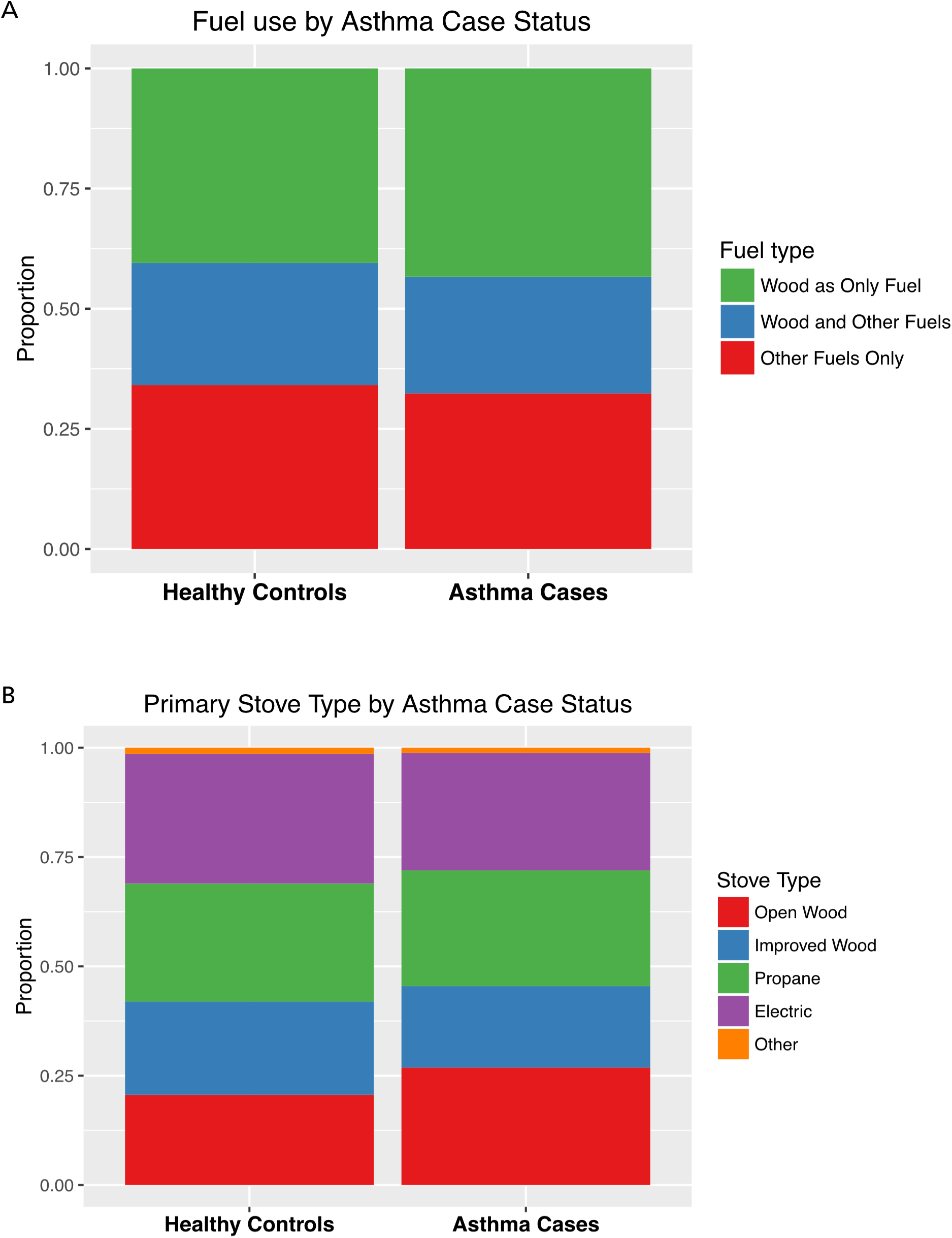
Unadjusted association between A) fuel use and asthma case status and B) primary stove type and asthma case status. While there was no significant difference in asthma case status by fuel type use, a larger proportion of cases with asthma used an open (unventilated) wood stove than controls.

After adjusting for age, sex, urbanization, maternal education, annual income, and household smokers, there was a significant association between use of open wood stove as main cooking source and asthma (OR = 1.8, 95% CI 1.2 – 2.8; p = 0.007), but not between the use of an improved (ventilated) wood stove and asthma (OR = 1.12, CI 0.7 – 1.8, p = 0.6) when compared to the reference group (electric stove) (Figure 3). Because there was no significant difference in the odds of asthma between electric, propane, and improved (ventilated) wood stoves, these categories were collapsed into a category (“other stoves”). Exposure to an open wood stove was associated with an increased risk of asthma (OR = 1.55, 95% CI 1.08 - 2.22, p = 0.02) compared to all other stoves. In the multivariate model, we found no association between degree of urbanization and asthma (ANOVA p-value = 0.2). Compared to participants living in rural environments, those living in small towns had an odds ratio for asthma of 0.72 (95% CI = 0.38 to 1.3, p = 0.3), while those living in an urban environment had an odds ratio of asthma of 0.61 (95% CI 0.33 to 1.1, p = 0.1).

**Figure 3.**
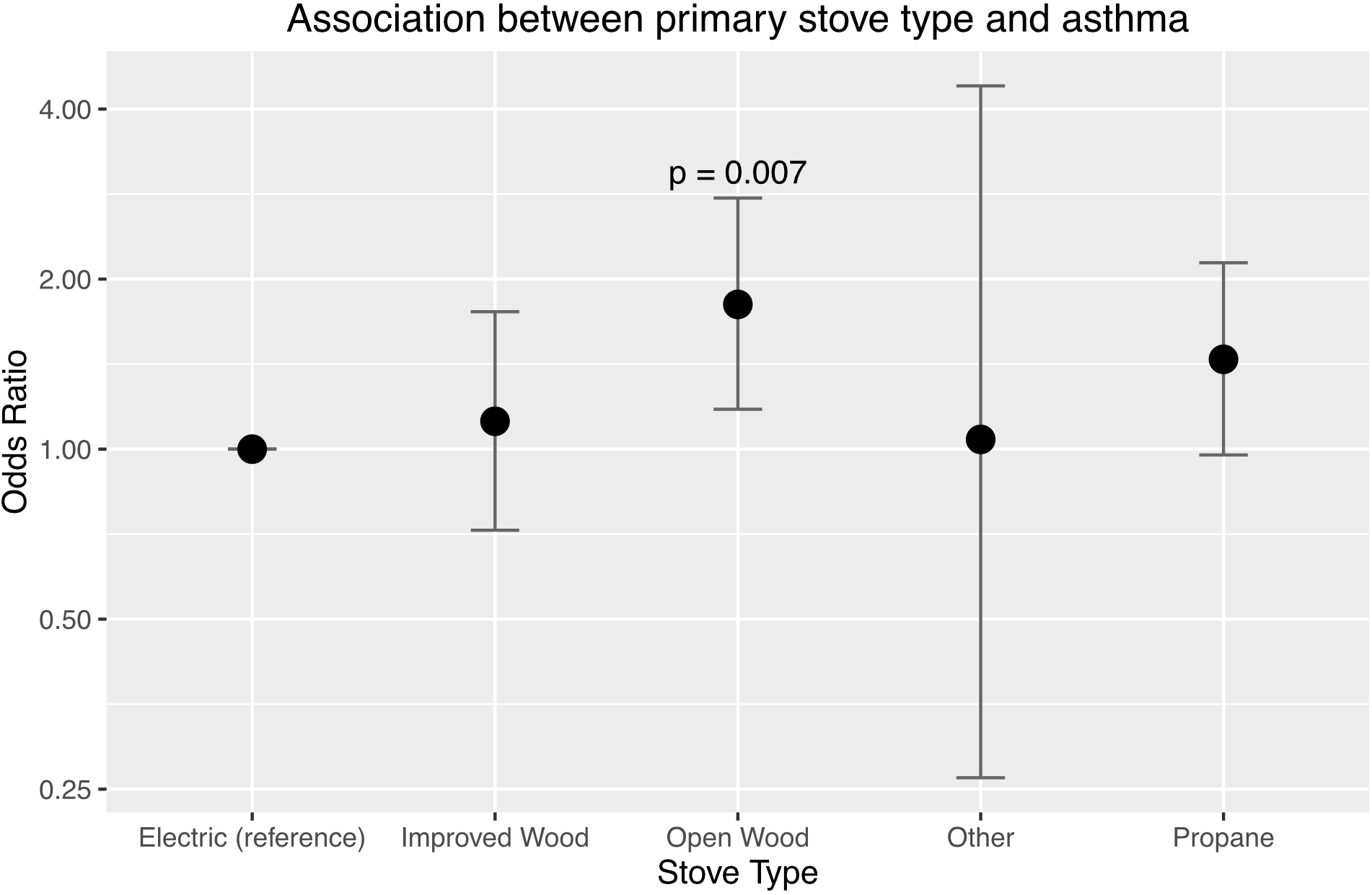
Adjusted association between primary stove type and asthma. While there is no association between fuel type and asthma, there was a significant association between the use of open wood stoves as a primary stove type and asthma (compared to electric stoves, the reference group). There was no significant association between improved wood stoves and asthma.

There was no association between increasing use of wood as a fuel and asthma diagnosis (OR 1.2 per change in category between exclusive use of other (non-wood) fuels, use of both wood and other fuels, or use of wood as sole fuel, p = 0.23). Collapsing the first two categories of wood use into two categories also showed no significant association between wood use and asthma diagnosis (OR = 1.18, p = 0.4).

The use of wood as fuel, however, was associated with asthma symptoms among those with an asthma diagnosis. There was a significant association between any use of wood as fuel and increased category of daytime symptom frequency (OR = 1.46, CI = 1.01-2.58, p =0.046) and nocturnal symptom frequency (OR = 2.51, CI = 1.04-2.62, p = 0.04), compared to the group that used only (non-wood) sources of fuel. There were no significant associations between use of wood as fuel and activity limitations or rescue inhaler use. Although participants with asthma who used wood had a score of 0.52 units lower on a modified version of the Asthma Control Test, the results were not statistically significant (CI = 1.23 lower to 0.19 higher, p = 0.15). We did not find any association between the type of fuel used or the type of stove used and pulmonary function (FEV_1_), adjusted for age, sex, height2, urbanization, maternal education, annual income, presence of household smokers, and inhaled corticosteroid use or FEV_1_ as a percent predicted. Specifically, use of an unvented wood stove was associated with a non-significant 47 mL increase in FEV_1_ compared to participants who used an electric stove (95% CI -86 mL to 179 mL, p = 0.5), which corresponds to a difference of 0.6 percent predicted. Likewise, use of an open wood stove was associated with a difference in FEV_1_/FVC ratio of 1.1% (95% CI -1.6% to 3.7%, p = 0.4). A similar effect is seen among controls without asthma, in which use of an open wood stove was associated with a 43 mL decrease in FEV_1_ compared to participants who used an electric stove (95% CI -178 mL to 93 mL, p = 0.5), corresponding to a difference of -0.3 percent predicted, and a 0.13% increase in the FEV_1_/FVC ratio (95% CI = -0.9% to 0.4%, p = 0.3).

Among healthy controls, use of wood as fuel was significantly associated with increased odds of non-asthmatic atopy (OR = 1.94, CI = 1.14 – 3.33, p = 0.015), defined as having allergic rhinitis, atopic dermatitis, or both. We did not find a similar association between the use of an open wood stove and atopy amongst cases with asthma. However, 89.6% of the participants with asthma had concomitant allergic symptoms so the study is underpowered. Among participants without asthma, there was also a significant association between use of wood as fuel and increased odds of chronic cough (OR = 2.22, CI = 1.09 – 4.88, p = 0.04), defined as more than 2 weeks of cough in the previous 12 months in the absence of an upper respiratory infection.

## Discussion

The results from this case-control study show that use of an open wood stove, but not a ventilated stove, is significantly associated with increased odds of asthma, and use of wood as a cooking fuel was associated with increased daytime respiratory symptoms and nocturnal respiratory symptoms among participants with asthma. These findings support previous findings from the International Study of Asthma and Allergies in Childhood (ISAAC), a large, multinational, cross-sectional study, which showed that use of open fires for cooking was associated with increased wheezing, reported asthma, and eczema among children aged 6-7 years and 13-14 years^18^. While in the ISAAC study did not identify a significant number of respondents with ventilated cooking stoves, in the present study, 19.8% of participants used an improved (ventilated) wood stove as their primary stove type, compared to 24.2% who had open wood stove. This is partly due to the efforts of a government program to distribute the “ecofogón”, a ventilated wood-fired stove with improved combustion efficiency and a built-in chimney with an external exhaust. Participants using this type of stove did not have increased odds of asthma compared to participants using an electric stove. In contrast, participants using an unventilated wood stove had 1.8 times higher odds of asthma than participants using an electric stove. This suggests that distribution of improved wood fired stoves like the ecofogón may reduce the risk of asthma imparted by unventilated wood stove use.

Participants with asthma in the present study were diagnosed by physician, in addition to meeting additional inclusion criteria of active treatment for asthma or current symptoms of asthma, which would likely reduce the potential for misclassification compared to the study by Wong et al. Nonetheless, the odds ratio for asthma (1.8, 95% CI 1.2 – 2.8) identified in this study are similar to those identified in the ISAAC study (1.78, 95% CI 1.51 – 2.10 for children 6-7 and 1.20, 95% CI 1.06 – 1.37 for children 13-14). The results of this paper thus provide further evidence that exposure to wood smoke may cause current clinical asthma.

We did not find evidence that the use of wood as a cooking fuel was associated with a higher risk of asthma, even though the use of an unimproved wood stove was a risk for asthma. This may reflect that in our study, participants who used wood as cooking fuel included both participants who used an open wood stoves as their main stove and participants who used an improved wood stove with ventilation as their main stove, and because a significant proportion of participants (24.8%) used multiple fuels for cooking (typically one wood stove and either a propane or electric stove). Since our findings showed that the risk of asthma is comparable between participants who had an improved (vented) wood stove and those who used non-wood fueled stoves (primarily gas or electric), combining both open stoves and vented stoves into one group masked the effect of wood smoke from using an open wood stove on asthma.

Indeed, ventilated wood stoves carried comparable odds for asthma to the referent (electric stove) category, and had meaningful, though not statistically significant, lower odds of asthma than open wood stoves (OR = 0.65, 95% CI = 0.41-1.02, p = 0.06).

As expected, there was a strong correlation between use of wood as cooking fuel and annual income as well as other measures of socioeconomic status such as maternal education and urban versus rural environment; participants with lower SES were more likely to use wood as a cooking fuel [see figure 1].

Neither use of wood as a cooking fuel nor stove type appeared to be associated with changes in pulmonary function as measured by FEV_1_/FVC, FEV_1_, or FEV_1_ as a percent of predicted, a finding consistent in both cases and controls. Although exposure to smoke from solid fuels used for cooking and heating is the second leading cause of chronic obstructive pulmonary disease in adults over 40, our population had a median age of 21, and inclusion in the study was limited to participants under 40 who had not been diagnosed with any respiratory diseases other than asthma, so as to minimize the possibility of including participants with COPD. Thus, although long-term exposure to wood smoke is known to cause obstructive lung disease, it is possible that our participants had not been exposed for long enough to cause a detectable change in lung function.

We found that both the use of wood as cooking fuel and use of unvented wood stove were associated with doubling of the odds of atopy among the non-asthmatic controls (OR = 1.94, CI = 1.14-3.33, p = 0.015) and (OR = 2.01, CI = 1.11-3.84, p = 0.026), respectively. Increasing air pollution has been associated with increasing prevalence of allergic diseases^35–38^. We hypothesize that household air pollution from wood smoke may lead to atopic sensitization via Th_2_-mediated IgE and cytokine production, resulting in atopic symptoms. This may represent one plausible mechanism by which exposure to wood smoke may lead to asthma.

We also observed that non-asthmatic controls who use wood as cooking fuel had a significantly increased risk of having chronic cough not associated with upper respiratory infections. This may be due to the irritant effects of wood smoke on respiratory epithelium.^32^ Healthy volunteers exposed to wood smoke in a human exposure chamber induced increases in airway epithelial permeability and fraction of exhaled nitric oxide (F_E_NO), which has been used as a biomarker that is elevated in patients with asthma and suggests evidence of eosinophilic inflammation. This points to the possibility that wood smoke may cause oxidative damage and allergic inflammation to the airways, and can lead to asthma in genetically susceptible individuals. Alternatively, as wood smoke exposure is associated with COPD, it may represent patients with undiagnosed COPD and preserved lung function (GOLD stage 0^33^ or COPD foundation stage 0^34^). Thus, though we did not see an association between wood stove use and lung function decline, even in our age range, wood stove use may place individuals at risk for activity limitation, COPD exacerbations, and airway disease.

Because the study is a case control study, it is susceptible to selection bias, though care has been taken to match the cases and controls with respect to age, sex, socioeconomic status, and geography. In addition, because all data were gathered in a form of a questionnaire, the study is susceptible to recall bias, and the study could have been biased by non-responses, even though there is no reason to believe that misclassification would be differential with respect to exposure or outcome, or that the missing data would not be missing completely at random. Thus, we would expect the results to have been biased towards the null. Despite these limitations, the results of this study provide more evidence for the potentially damaging effects of wood smoke on asthma and respiratory symptoms and encourage more controlled interventional trials to evaluate the effect of improved wood stoves or electric stoves on risk of asthma and other obstructive lung diseases. The fact that the risk is associated with unvented wood stoves but not improved (ventilated) stoves supports further implementation of these types of stoves in areas where electricity or other cleaner stoves cannot be introduced.

Extensive characterization of the relationship between use of wood stoves, stove type, and household measurements of air pollution such as CO, NO, and particulate matter (PM_10_ and PM _2.5_) in the field would strengthen the link between unimproved wood stoves and exposure. Further linking levels of household air pollution to personal exposure using biomarkers of wood smoke exposure (such as urinary methoxyphenols), as well as airway inflammation and allergic response (such as IgE, IL5 and IL13) among subjects with and without asthma who use wood stoves would further elucidate the mechanism by which wood smoke may cause asthma in genetically susceptible individuals. Moreover, testing gene by environment associations with asthma, atopy, and related traits in the context of wood smoke exposure, specifically single nucleotide polymorphisms in antioxidant genes that play a role in airway oxidative damage^40^, may provide further mechanistic insight into how gene-environmental interactions may drive asthma pathogenesis and identify individuals who are genetically susceptible to the damaging effects of wood smoke.

## Conclusions

Our data suggests that exposure to wood smoke from unvented wood stoves is a significant risk factor for asthma and respiratory symptoms. Wood smoke is associated with increased odds of atopy and chronic cough not related to upper respiratory infections among controls without asthma. Wood smoke may play an important role in allergic sensitization, asthma pathogenesis, and development of obstructive lung disease in genetically susceptible individuals. Replacing open wood stoves with improved wood stoves with ventilation or even cleaner gas or electric stoves may significantly improve symptoms in asthmatic individuals and decrease the prevalence of asthma and other obstructive lung diseases in the developing world.

## Acknowledgements

The authors would like to thank the participants and their families for their participation in this study. We would also like to thank Hospital Santo Hermano Pedro in Catacamas, Honduras and Hospital San Francisco in Juticalpa, Honduras and the numerous healthcare providers at the hospitals who facilitated the study. We are also deeply indebted to the Universidad Católica de Honduras, Dr. Jorge Humberto Melendez Bardales, Dean of the School of Medicine, and Dra. Shirley Andino Moya, Academic Director at the School of Medicine, and the many students who were instrumental in patient recruitment, specifically Dra. Vilma Georgina Rodriguez Bueso, Dra. Claudia Elizabeth Chaparro Ramos, Dra. Nitza Ivonne Valladares Hernandez, Dra. Angela Mariela Sosa Ramos, Eduardo Alberto Piloña Flores, Daniel Eduardo Andino Alvarado, Valeria Margarita Sierra Roca, Andrea Poleth Castellanos Ulloa, Aby Sabina Argueta Matute, Irma Cristina Argueta Matute, Gustavo Adolfo Flores Fortin. Computations in this manuscript were performed using the UCSF Biostatistics High Performance Computing System.

## Sources of Funding

This research was supported by the NHLBI (K23HL111636) and NCATS (KL2TR000143), within the National Institutes of Health. JMG was supported in part by NIH Training Grant T32 (T32GM007546) as well as the Hewett Fellowship; NT was supported in part by an institutional training grant from the NIGMS (T32-GM007546) and career development awards from the NHLBI (K12-HL119997 and K23-HL125551), Parker B. Francis Fellowship Program, and the American Thoracic Society. EGB was supported in part by the Sandler Family Foundation, the American Asthma Foundation, the RWJF Amos Medical Faculty Development Program, National Institutes of Health 1R01HL117004, R01Hl128439, R01 ES015794, R21ES24844, 1P60 MD006902, U54MD009523, 1R01MD010443 and the Tobacco-Related Disease Research Program under Award Number 24RT-0025. This publication was supported by various institutes within the National Institutes of Health. Its contents are solely the responsibility of the authors and do not necessarily represent the official views of the NIH.

